# Scm3 interacts with the N-terminal tail of Cse4 to regulate kinetochore assembly in budding yeast

**DOI:** 10.1101/2025.10.20.683541

**Authors:** Prakhar Agarwal, Anushka Alekar, Shubhomita Mallick, Jaladhi Shah, Munira Basrai, Santanu Kumar Ghosh

## Abstract

The kinetochore is a multiprotein complex formed at the centromeres and is essential for the faithful chromosome segregation. In most of the organisms the kinetochore is assembled on a specialized centromeric nucleosome where histone H3 is replaced by a variant, named CENP-A. In budding yeast, the CENP-A homolog, Cse4 is recruited to the centromeric nucleosome through an interaction between its C-terminal domain and a specific chaperone, named Scm3. Interestingly, following Cse4 recruitment during S phase, Scm3 persists at the centromeres throughout the cell cycle. Recent in vitro studies have reported that Scm3 also interacts with N-terminal of Cse4 (N-C) that in turn facilitates a better interaction of Ame1-Okp1 (AO) of COMA subcomplex with N-C, which promotes kinetochore assembly. In this work, using genetic and biochemical assays we provide in vivo evidence of the interaction between Scm3 and N-C. Additionally, by artificially tethering Scm3, we show that their association has the potential to stabilize a missegregating chromosome with inactive centromere, perhaps by forming an active kinetochore at an ectopic site. We propose that at the centromeres, Scm3 has two jobs in tandem - Cse4 deposition and stabilization of N-C, which together culminates into proper kinetochore assembly. This work has clinical significance as both CENP-A and its chaperone are known to be upregulated in human cells under disease states which can predispose the cells to aneuploidy due to increased chances of forming ectopic kinetochores.

## Introduction

The centromere is a unique chromosomal locus on DNA that recruits the kinetochore complex for faithful chromosome segregation (Verdaasdonk and Bloom, 2011). In most of the organisms, the centromere is epigenetically determined by the presence of a histone H3 variant called CENP-A (Centromere Protein A) (Earnshaw and Rothfield, 1985). The formation of active functional centromeres in these organisms relies on the replacement of both the copies of H3 by CENP-A at the centromeric nucleosomes rather than on the centromeric DNA sequence per se. Given the importance of CENP-A on centromere formation, its deposition at the centromeres is highly regulated (Da Rosa et al., 2011; Hildebrand and Biggins, 2016; Renaud-Pageot et al., 2022). Any misregulation of intracellular CENP-A level can lead to its deposition at the non-CEN loci, resulting in the formation of ectopic centromeres generating aneuploidy based diseases, including cancers (Li et al., 2011; Ma et al., 2003; McGovern et al., 2012; Wu et al., 2012). The CENP-A containing nucleosome promotes the formation of kinetochore as it differs from the H3-nucleosome in a few different ways. CENP-A has an elongated loop L1 (Phe78-Phe84) at its N terminal, which contains two additional amino acids, Arg80 and Gly81, compared to H3. These residues are located at the tip of the L1 loop that protrudes out from the nucleosome (Tachiwana et al., 2011) and are key for the stable retention of CENP-A at the centromere and also for the binding of other centromeric proteins. Additionally, the presence of a shorter αN helix (48 to 55 residues) in CENP-A compared to H3, allows the DNA at these sites to adopt specific conformations important for centromere protein binding (Quénet and Dalal, 2012; Tachiwana et al., 2011). Since the C terminal tail (138 to 143 residues) of CENP-A is disordered, this enables the binding of the kinetochore protein, CENP-C (Tachiwana et al., 2012). It has been shown that the CENP-A nucleosome occupies a shorter stretch of DNA of approximately 121-125 bp compared to H3 nucleosome that protects 147 bp (Tachiwana et al., 2011). Consequently, the DNA at the entry and exit points of the CENP-A nucleosome is free and is probably important for the interaction of another kinetochore protein CENP-B. While the C-terminal histone fold domain (HFD) of CENP-A (118 to 134 residues) is highly conserved across eukaryotes, the N terminal domain (1 to 74 residues) of which the region 1 to 38 residues is disordered, differs across organisms (Tachiwana et al., 2011).

Cse4, the CENP-A in budding yeast, has an extra-long N terminal domain of 135 residues, henceforth called N-C (N-terminal of Cse4), which has been shown to be important for its centromeric localization and promoting kinetochore assembly (Chen et al., 2000; Keith et al., 1999). For instance, the N-C undergoes several post-translational modifications that contribute to Cse4 recruitment at the centromeres (Boeckmann et al., 2013; Hoffmann et al., 2018; Samel et al., 2012); it regulates the level of Cse4 through proteolysis of Cse4 by interacting with Psh1, the major ubiquitin ligase (Au et al., 2013; Cheng et al., 2017). The N-C also promotes kinetochore assembly through an association with the COMA sub-complex of the Ctf19 complex of budding yeast kinetochore (Anedchenko et al., 2019; Fischböck-Halwachs et al., 2019). Moreover, recently the N-C has also been implicated in imparting stability of Cse4 at the centromeres (Popchock et al.,2025).

Systematic mutagenesis studies have shown that a small region from 28 to 60 residues within N-C, characterized as the Essential N-terminal Domain (END), is essential for Cse4 function (Chen et al., 2000). Further, the residues 34 to 46 within END were found critical for the interaction with the essential subunits of the COMA sub-complex, the Ame1 and Okp1 (AO) (Fischböck-Halwachs et al., 2019). Mutations either in the END domain or in the END domain interacting residues in AO have a direct impact on cell viability (Deng et al., 2023). This suggests that AO forms a strong connection between the centromeres and the microtubules where N-C plays a significant role (Hamilton et al., 2020, Deng et al., 2023).

Post-translational modifications of the N-C are essential for its interaction with the kinetochore and other proteins that contribute to its stability and its deposition at the centromeres. For instance, methylation and acetylation of the R37 and K49 residues, respectively, regulate the interaction of Cse4 with the AO (Fischböck-Halwachs et al., 2019; Samel et al., 2012). Sumoylation of K65 significantly negates ubiquitin mediated proteolysis by Psh1 and thus regulates the stability of Cse4 (Ohkuni et al., 2018). Additionally, phosphorylation of S33 facilitates the deposition of Cse4 at the centromeres, perhaps by promoting interaction with its chaperone, Scm3 (Hoffmann et al., 2018).

Scm3 has been characterized as a Cse4 chaperone, since it promotes deposition of Cse4 at the centromeres through physical interaction with Cse4 and the inner kinetochore protein, Ndc10, and it was found at the centromeres (Camahort et al., 2007, Stoler et al., 2007). It has been demonstrated that both the absence and overexpression of Scm3 leads to reduced localization of Cse4 at the centromere (Mishra et al., 2011). Although the deposition of Cse4 at the centromere occurs only during a short window in the S phase (Pearson et al., 2004; Shivaraju et al., 2012), Scm3 is retained at the centromeres throughout the cell cycle (Xiao et al., 2011) indicating that it might have additional functions at the centromeres besides the deposition of the Cse4. While Scm3 deposits Cse4 to the centromere through an interaction with the HFD of Cse4, recent in vitro studies have shown that Scm3 and the N-C also physically interact (Popchock et al., 2025) perhaps promoted by their intrinsically disordered entities (Shukla et al., 2024), which also provides a hint to the additional functions of Scm3 at the centromeres. It is suggested that the Scm3-N-C interaction stabilizes the two proteins, lending a more open structure to Scm3 for it to interact with the kinetochore proteins AO. The Scm3-AO interaction then, in turn, promotes a stronger interaction between N-C and AO required for the kinetochore assembly (Shukla et al., 2024).

Given the above evidence of Scm3 and N-C interaction from in vitro studies, we hypothesized that an optimum level of their in vivo interaction is crucial for proper kinetochore assembly and faithful chromosome segregation and wished to test this using genetic and biochemical assays. In this work, we report that the misregulated levels of N-C cause chromosome loss phenotype, which becomes aggravated in the presence of overexpression of Scm3. The interaction between the two proteins titrates away essential components of COMA complex from the kinetochore. This results in a segregation incompetent kinetochore and eventually leads to cell death. This work suggests that the interaction between Scm3 and N-C strengthens the COMA complex binding at the kinetochore and thus provides first-hand in vivo evidence of significance of this interaction. This work demonstrates an additional role of Scm3 at the centromeres which can be tested for the homologous protein HJURP in metazoans.

## Results

### Overexpression of N-C causes synthetic sick phenotype in the kinetochore mutants

Given the crucial role of the N-C in chromosome segregation in budding yeast (Chen et al., 2000; Keith et al., 1999), it is important to understand how it confers that role. Since the END domain of N-C interacts with the COMA proteins to initiate kinetochore assembly (Anedchenko et al., 2019; Fischböck-Halwachs et al., 2019), it is expected that upon over expression of N-C, the excess N-C would sequester the COMA proteins and lead to weak kinetochore formation at the centromeres causing poor cell viability. To examine this, we overexpressed N-C using galactose inducible promoter from a multi-copy episomal vector, named *Gal N-C* (pMB1236, from Basrai lab). Wild type cells transformed with *Gal N-C* or an empty vector were grown till mid-log, and serial dilutions of the strains were spotted on dextrose or galactose containing plates. However, we did not observe any growth defect of wild type cells overexpressing N-C (Figure 1A). This could be due to the ability of wild type cells to negate the effects of overexpressed N-C and still hold a segregation competent kinetochore. However, we presumed that the non-essential kinetochore protein mutants may not be able to negate it. Therefore, we wished to examine how such mutants behave in response to higher levels of N-C. We used *ctf19Δ* and *mcm21Δ* mutants as the representatives from the COMA sub-complex and used the *ctf3Δ* mutant as a representative from the adjacent Ctf3 sub-complex. These mutant cells were also transformed with *Gal N-C* or empty vector and subjected to spotting assay. We observed a drastic drop in cell viability in the COMA sub-complex kinetochore mutants (*ctf19Δ* and *mcm21Δ)*, but *ctf3Δ* cells did not show such sensitivity (Figure 1A). We reasoned that in the absence of Ctf19 or Mcm21, when the COMA sub-complex is already compromised, the sequestration of AO by excess N-C perhaps further weakened the COMA sub-complex at the centromeres, causing cell death.

**Figure 1:**
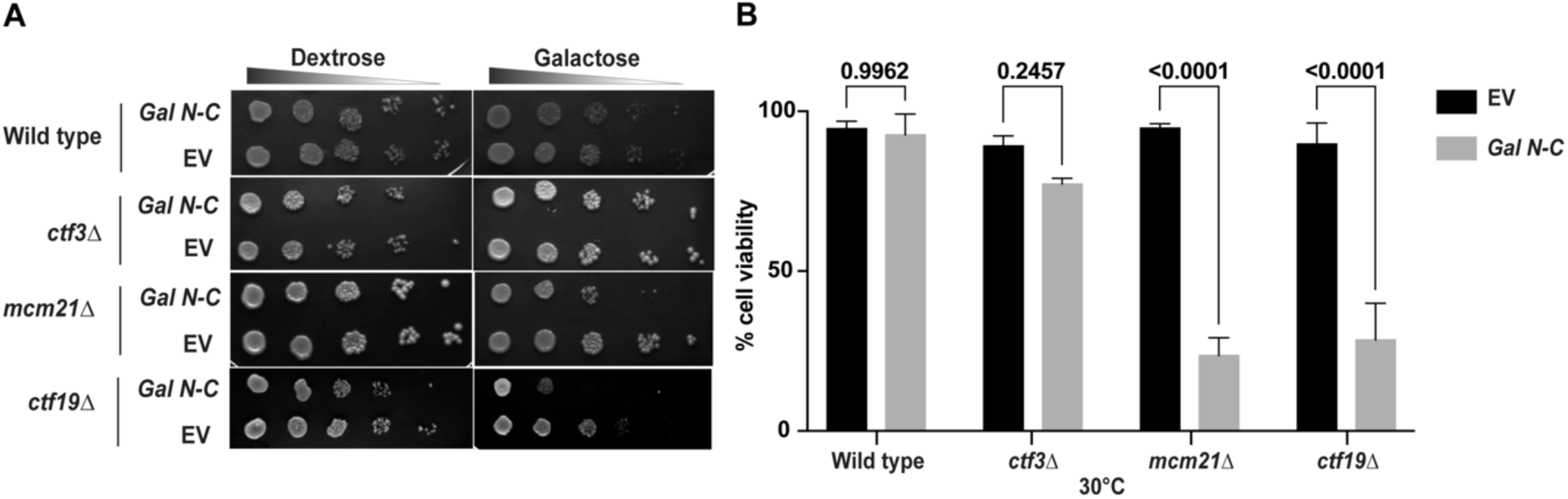
Overexpression of N-C leads to loss of cell viability in kinetochore mutants. **(A)** Wild-type, *ctf3Δ, mcm21Δ, and ctf19Δ* cells harboring *Gal N-C* or empty vector (EV) were grown till mid-log before they were serially diluted and spotted on the indicated plates. The plates were incubated at 30°C for 24-48 hrs before imaging. **(B)** An equal number of cells from (A), were spread on plates containing either dextrose or galactose. From the number of colonies that appeared after incubation at 30°C for 2-3 days, the cell viability was calculated (see materials and methods) and plotted. Error bars represent the standard deviation from the mean values obtained from three independent experiments. For each experiment, at least 100 colonies were counted. The statistical significance *p* value was calculated using an unpaired two-tailed student’s t-test.

To quantify the extent of sensitivity towards overexpression of N-C, we performed a CFU count assay where wild type, *ctf19Δ, mcm21Δ,* and *ctf3Δ* strains harboring either *Gal N-C* or empty vector were grown till mid log and an equal number of cells were plated on dextrose and galactose plates. The number of colonies formed after 24 or 36 hrs on dextrose or galactose plates, respectively were counted, and the cell viability was quantified. We observed a statistically significant drop in cell viability in *ctf19Δ* and *mcm21Δ* strains, while wild type and *ctf3Δ* strains did not show such sensitivity (Figure 1B).

We further tested the ability of the kinetochore mutants to withstand the detrimental effect of overexpression of N-C by imposing a temperature stress by growing the cells as in Figure 1A but at lower temperatures such as 16°C and 11°C. We observed that while the wild type showed very little sensitivity to overexpressed N-C, *ctf3Δ* showed significant sensitivity at both 11°C and 16°C (Figure S1). As expected, the effect of excess N-C on *ctf19Δ* and *mcm21Δ* cells was found to be more severe at these low temperatures compared to 30°C. We conclude that the excess N-C in the kinetochore mutants makes them synthetically lethal, probably by sequestering the kinetochore proteins, reducing their normal stoichiometry at the centromeres required for proper kinetochore formation.

### Scm3 and N-C interact causing an aggravated sick phenotype in the kinetochore mutants

Having observed that overexpression of N-C weakens the kinetochore, causing loss of cell viability, we aimed to test if this phenotype becomes aggravated in the presence of excess Scm3. This aim is based on the recent findings from studies reporting that Scm3 is important to augment interaction between N-C and the kinetochore proteins, AO (Shukla et al., 2024, Popchock et al., 2025). From this, we hypothesize that excess Scm3 can lead to more efficient titration out of the kinetochore proteins by overexpressed N-C, causing further weakening of the kinetochore as well as a drop in cell viability. To test this, we overexpressed both N-C and Scm3 from the *GAL* promoter in the same cells of wild type and *ctf3Δ* and *ctf19Δ* strains. The cells pregrown in dextrose or galactose were spotted on the dextrose or galactose plates, respectively, along with the appropriate controls (Figure 2A). The wild type cells hardly showed any sensitivity upon over expression of either N-C or Scm3 or both. Although the CFU count assay, performed as described above, showed a significant drop in cell viability when the both the proteins were overexpressed (Figure 2B, wild type). On the other hand, both the mutants *ctf3Δ* and *ctf19Δ* showed considerable growth defects when either N-C or Scm3 was overexpressed, and the defect was found to be very severe when both the proteins were overexpressed (Figure 2A). This was also reflected in the CFU count assay and more defect than the spotting assay was detected in *ctf19Δ* compared to *ctf3Δ* cells even when overexpressing either N-C or Scm3 (Figure 2B). As expected, the growth defects were found proportionately aggravated at lower temperature (Figure S2A, B). Taken together, we provide in vivo evidence of Scm3 interacting with N-C from our findings that overexpression of Scm3 aggravates the growth defects of the kinetochore mutants overexpressing N-C, which possibly happens due to more efficient siphoning out of the kinetochore proteins from the centromeres by excess N-C in presence of excess Scm3; this supports earlier claims by the in vitro studies (Shukla et al., 2024, Popchock et al., 2025).

**Figure 2:**
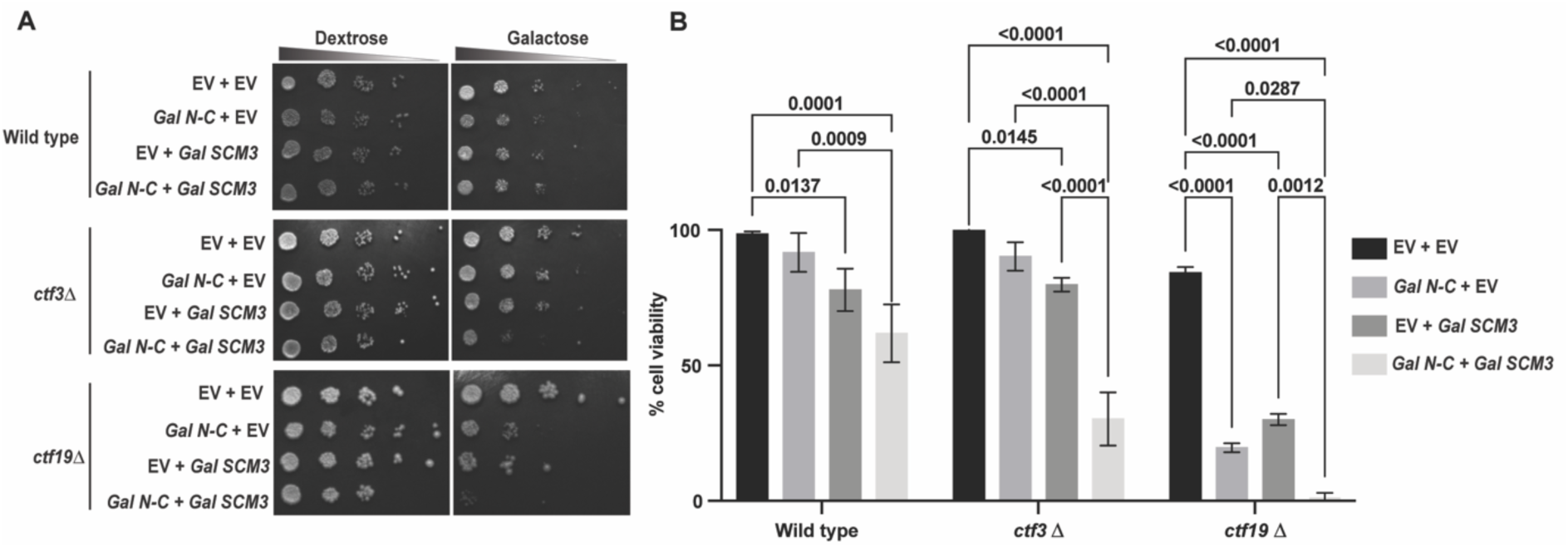
Overexpression of both N-C and Scm3 causes synthetic sick phenotype. **(A)** Wild type, *ctf3Δ,* and *ctf19Δ* cells harboring both *Gal N-C* and *Gal SCM3,* or corresponding empty vector (EV) in different combinations, were grown till mid-log before they were serially diluted and spotted on the indicated plates. The plates were incubated at 30°C for 24-48 hrs before imaging. **(B)** An equal number of cells from (A), were spread on plates containing either dextrose or galactose. From the number of colonies that appeared after incubation at 30°C for 2-3 days, the cell viability was calculated (see materials and methods) and plotted. Error bars represent the standard deviation from the mean values obtained from three independent experiments. For each experiment, at least 100 colonies were counted. The statistical significance *p* value was calculated using a one-way ANOVA test.

### Overexpression of Scm3 and N-C reduces enrichment of kinetochore proteins at the centromeres

From the above results, we assumed that an excess amount of Scm3 and N-C, through their interaction with the kinetochore proteins (AO in particular), reduces the level of these proteins at the centromeres, causing sick growth phenotype of the *ctf3Δ* or *ctf19Δ* cells. Therefore, we wished to test the level of Ame1 at the centromeres in *ctf19Δ* cells overexpressing N-C and Scm3 using cell biological and biochemical assays. We compared Ame1-13Myc localization at the centromeres in *ctf19Δ* cells harboring empty vector, overexpressing N-C or Scm3 or both N-C. To negate any effect due to variation in the pace of the cell cycle among the strains, we arrested the cells at the metaphase stage by using nocodazole and performed immunofluorescence assay to visualize Ame1-13Myc in such cells (Figure 3A). We presumed that proper localization of Ame1 at the centromeres would generate ‘clustered’ kinetochores, whereas any reduction in its level can produce ‘declustered’ or ‘weak signal’ category of kinetochores (Figure 3A). Notably, the ‘declustered’ phenotype may also arise due to perturbations in higher order assembly of the kinetochore without reducing the level of Ame1 at the centromeres. We could observe an increased percentage of cells with declustered kinetochores/weak signal, marked as Ame1-13Myc foci, in *ctf19Δ* cells overexpressing both N-C and Scm3 as compared to empty vector controls (75% vs 35%) or *ctf19Δ* cells overexpressing either N-C (75% vs. 57%) or Scm3 (75% vs. 52%) (Figure 3B). The sharp increase of the ‘declustered/weak signal’ category in the *ctf19Δ* cells overexpressing both N-C and Scm3 (Figure 3B) indicating loss of Ame1 from the centromeres, justifies their extensive loss of cell viability (Figure 2B). To rule out the possibility that the overall expression of Ame1 is perturbed in such cells, we performed a western blot analysis and found no significant difference in the protein levels among the strains (Figure S3).

**Figure 3:**
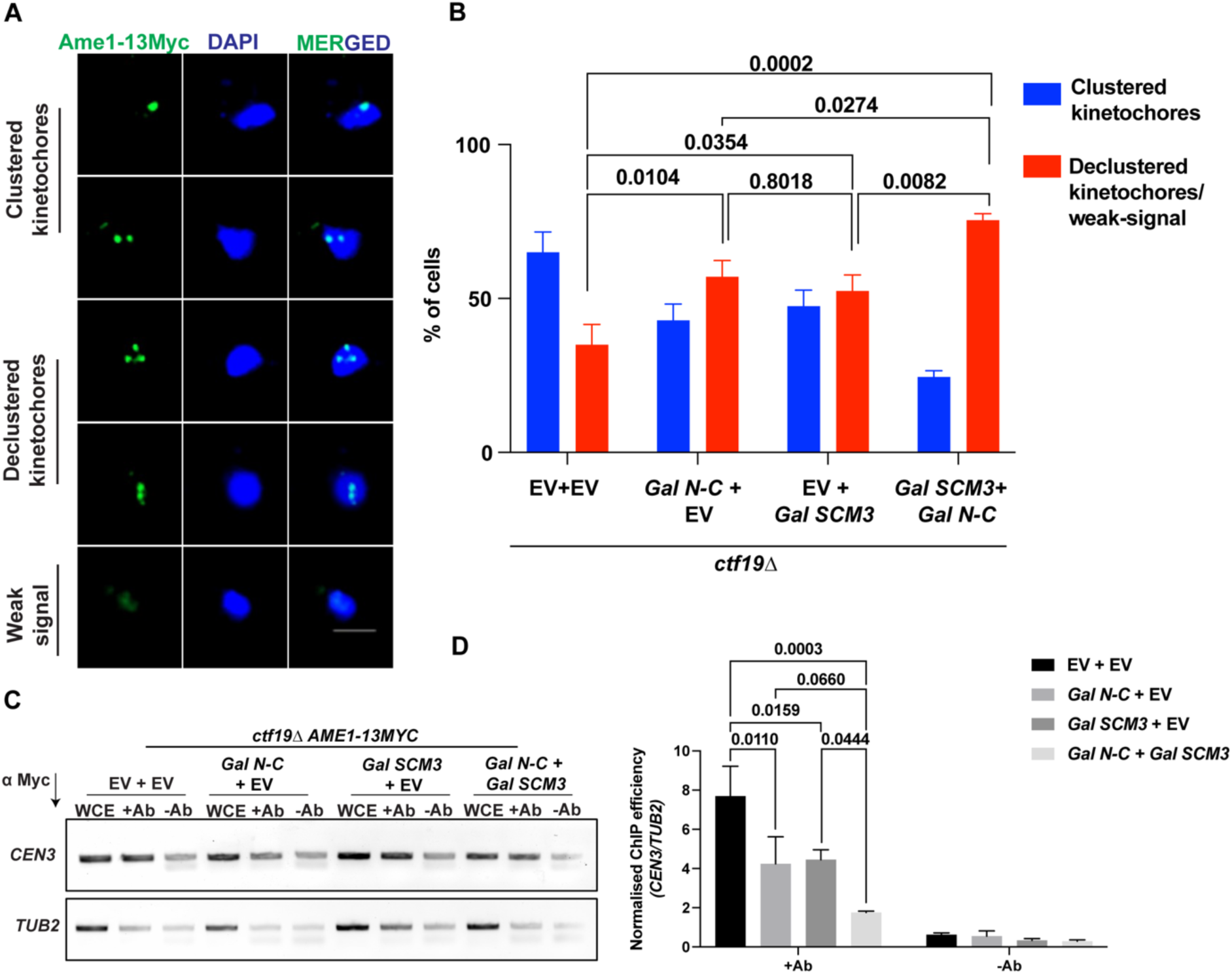
Overexpression of both N-C and Scm3 causes a defect in kinetochore organization. **(A)** The representative images of the immunofluorescence assay to visualize kinetochore protein Ame1 fused to 13Myc in *ctf19Δ* cells harboring indicated plasmids in metaphase arrested cells. Anti-Myc antibodies and DAPI staining were used to visualize Ame1-13Myc and chromatin, respectively. Scale bar = 2 µm **(B)** The percentage of different categories of the kinetochore phenotype, as mentioned in A, for the indicated strains. N = 100, Experimental replicates, n = 2. **(C–D**) ChIP-qPCR analysis showing the association of Ame1-13Myc with *CEN3* and *TUB2* (negative control) loci in the indicated strains. Anti-Myc antibodies (ab9106, Abcam) were used for immunoprecipitation in metaphase arrested cells. Error bars represent the standard deviation from the mean values obtained from two independent experiments. For each set in (B), at least 100 cells were counted. The statistical significance *p* value was calculated using a two-way ANOVA test for both (B) and (D).

To further validate that in *ctf19Δ* cells, the level of Ame1 at the centromere indeed goes down upon overexpression of N-C and Scm3, we examined the localization of Ame1-13Myc at the centromeres by chromatin immunoprecipitation (ChIP) assay in the metaphase arrested *ctf19Δ* cells in absence or presence of excess N-C+Scm3. We used anti-Myc antibodies to pull down Ame1-13Myc in the cells used in Figure 3A. While in the *ctf19Δ* cells harboring empty vectors, we could see an enrichment of Ame1-13Myc at the centromeres, there was a significant reduction in its enrichment in the *ctf19Δ* cells overexpressing N-C and Scm3 (Figure 3C-D) when normalised with respect to a negative locus, *TUB2*. With this, we conclude that Scm3 and N-C interact in vivo and their simultaneous overexpression titrates away key kinetochore proteins such as Ame1 from the centromeres and results in the loss of cell viability.

### Evidence of in vivo physical interaction between Scm3 and N-C

From the observed synthetic interactions between Scm3 and N-C and the reduction of Ame1 at the centromeres with overexpressed N-C and Scm3, we presumed that there is a physical interaction between Smc3 and N-C. This was also fueled by an earlier report of N-C and Scm3 physical interaction in vitro (Shukla et al., 2024). Since there is no direct in vivo evidence for the physical interaction we sought out to investigate this using the yeast two-hybrid assay (James et al., 1996) where the earlier shown interaction between Cse4 and Scm3 was taken as the positive control (Stoler et al., 2007). Full-length genes of Cse4, Scm3, and N-C were cloned in the requisite two-hybrid vectors, pGAD and pGBD expressing Gal4 transcriptional activation and DNA binding domains, respectively. We confirmed the functionality of AD-Cse4 or BD-Scm3 fusion proteins as they could rescue the growth in cells where endogenous Cse4 or Scm3, respectively was depleted by auxin-inducible degron (AID) system (Figure S4). The fusion proteins BD-Cse4 and AD-Sm3 were found non-functional (data not shown). We also could not verify the functionality of AD/BD-N-C fusion proteins due to lack of testable phenotypepGAD and pGBD with or without the genes of interest in different combinations were introduced into the PJ69-4A strain harboring the reporter genes *HIS3* and LacZ. Expression of *HIS3* on histidine drop out plate supplemented with 3-AT (to block basal level expression) and β-galactosidase activity due to LacZ expression were taken as the reporters for the positive interaction. To examine the expression of the *HIS3* reporter, PJ69-4A cells with different combinations of the plasmids were plated on SC-Leu-Trp-His media supplemented with indicated concentrations of 3AT (Figure 4A). As expected, we did not observe any growth for the negative control strains; minimal growth for BD-Scm3+AD, which was reported previously (Camahort et al., 2007) and strong growth for the positive control AD-Cse4+BD-Scm3 as reported earlier (Stoler et al., 2007) were observed. Importantly, we could also observe a significant growth of the cells harboring AD-N-C+BD-Scm3, which indicates that Scm3 indeed interacts with N-C in vivo. From the qualitative growth assay, it appears that the extent of interaction of Scm3 with N-C is lesser than its interaction with Cse4. To verify this quantitatively, we used the LacZ reporter assay and estimated the β-galactosidase activity in these cells and observed a similar extent of interactions (Figure 4B). Overall, from these assays we conclude that Scm3 physically interacts with N-C in vivo, however, not as strongly as it does with Cse4 (Figure 4C).

**Figure 4:**
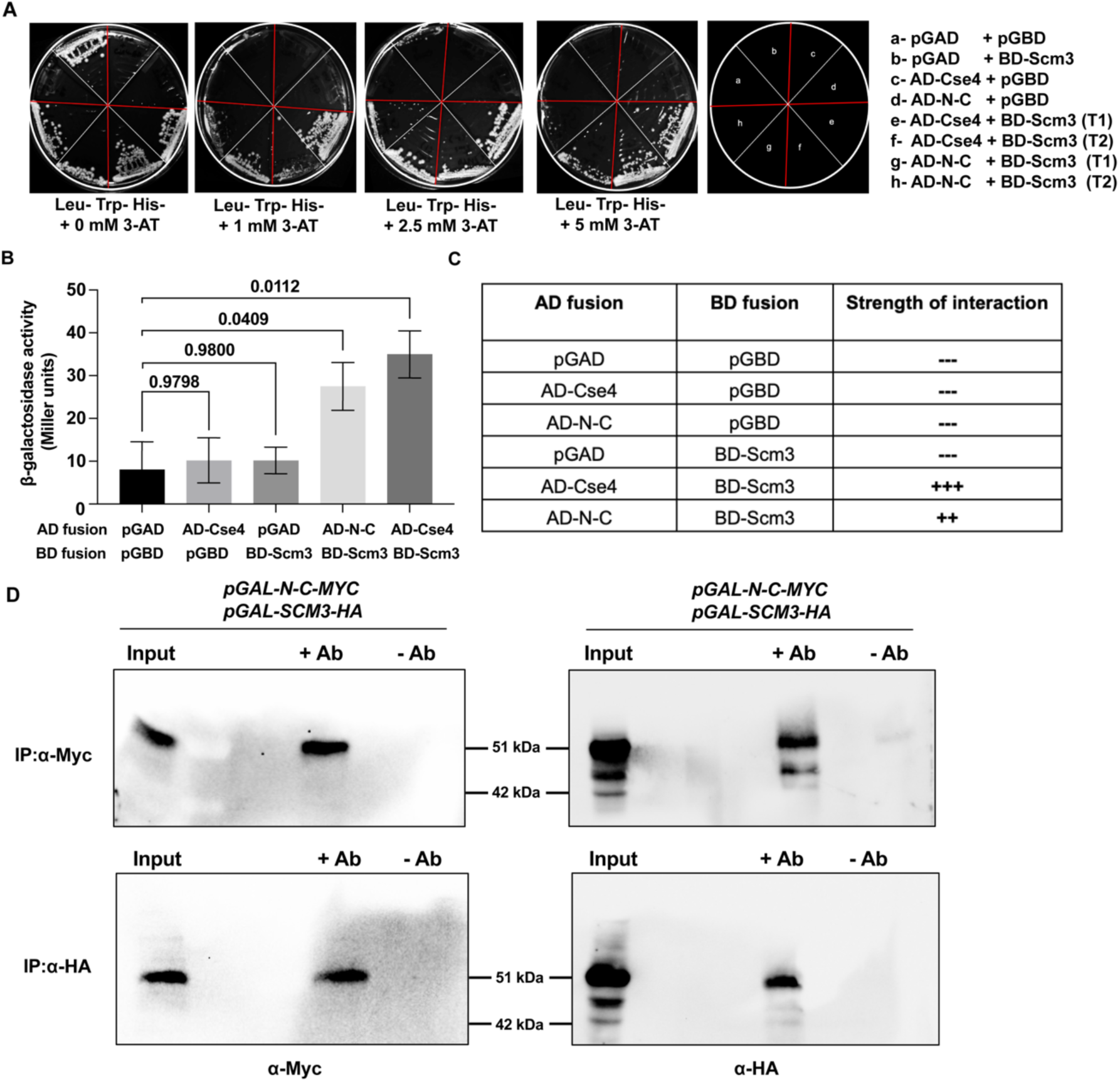
Physical interactions among Scm3, N-C, and Cse4. **(A)** PJ69-4A cells harboring indicated combinations of two hybrid plasmids were streaked on the indicated selection plates in the absence or presence of different concentrations of 3-AT **(B)** The same cells were used for β-galactosidase activity which is graphically represented. Error bars represent the standard deviation from the mean values obtained from two independent experiments. The statistical significance *p* value was calculated by the two-tailed student’s t-test for the mean. **(C)** The table summarises the strength of interaction obtained from different pairwise combinations. **(D)** Top panel, cell lysates from asynchronous cells expressing N-C-Myc and Scm3-HA under galactose inducible promoters were immunoprecipitated using the polyclonal anti-Myc (ab9106, Abcam) antibody. Scm3-HA and N-C-Myc were detected by immunoblotting using monoclonal anti-HA (3F10, Roche) and anti-Myc (9E10, Abcam) antibodies, respectively. Bottom panel, same as above except cell lysates were immunoprecipitated using the polyclonal anti-HA (ab9110, Abcam) antibody.

We further used a co-immunoprecipitation experiment to validate our yeast two-hybrid assay results on the physical interaction between Scm3 and N-C. Wild type cells co-expressing *SCM3-6HA* and *N-C-13MYC* from galactose inducible promoters borne in episomal vectors were used. Total protein was extracted from mid log grown cells; immunoprecipitation of N-C-13Myc pulled down Scm3-6HA and vice versa (Figure. 4D). These results reconfirm in vivo physical interaction between Scm3 and N-C.

### Ectopically targeted Scm3 interacts with N-C to improve chromosome segregation

Given the physical interaction between Scm3 and N-C and their unified ability to better interact with kinetochore proteins, we wanted to examine if ectopically targeted Scm3-N-C complex can drive chromosome segregation presumably by forming an active kinetochore. To elucidate this, we used a strain with a conditional *CEN* IV where under galactose the *CEN* IV becomes inactive and chromosome IV missegregates. We marked the chromosome IV close to *CEN* IV by GFP by integrating 226 copies of the Lac operators (LacO) at the *TRP1* locus and by expressing LacI-GFP (Figure 5A). The functioning of the strain was verified by monitoring the 1:1 equal segregation or 2:0 missegregation of chromosome IV (*CEN* IV-GFP) in anaphase cells as judged by divided DAPI signal (Figure 5B). While cells grown in dextrose showed a high frequency of equal segregation of chromosome IV (98%), in galactose it was much lower (2%), confirming the functionality of the strain as expected (Figure S5A). As per our hypothesis, the equal segregation of chromosome IV in galactose is expected to be rescued upon co-expression of Scm3-LacI and N-C together in these cells. This is because Scm3-LacI will be targeted to the *CEN* IV proximal LacO array and along with N-C, will promote formation of an active kinetochore there which cannot be inactivated by galactose. We first verified that Scm3-LacI fusion protein is functional as it is able to rescue the viability of *SCM3-AID* cells in presence of auxin (Figure S5B). As expected, we observed that Scm3-LacI and N-C together, but not individually, could rescue the equal segregation frequency of chromosome IV compared to GFP-LacI alone (Figure 5C). However, the extent of rescue was not as high as when Ask1-LacI was expressed, which was shown earlier to induce kinetochore formation (Lacefield et al., 2009). Nevertheless, our results indicate that ectopic targeting of Scm3 on a missegregating chromosome, along with its interaction with N-C, can alleviate the missegregation frequency possibly due to formation of an active kinetochore at that ectopic location.

**Figure 5:**
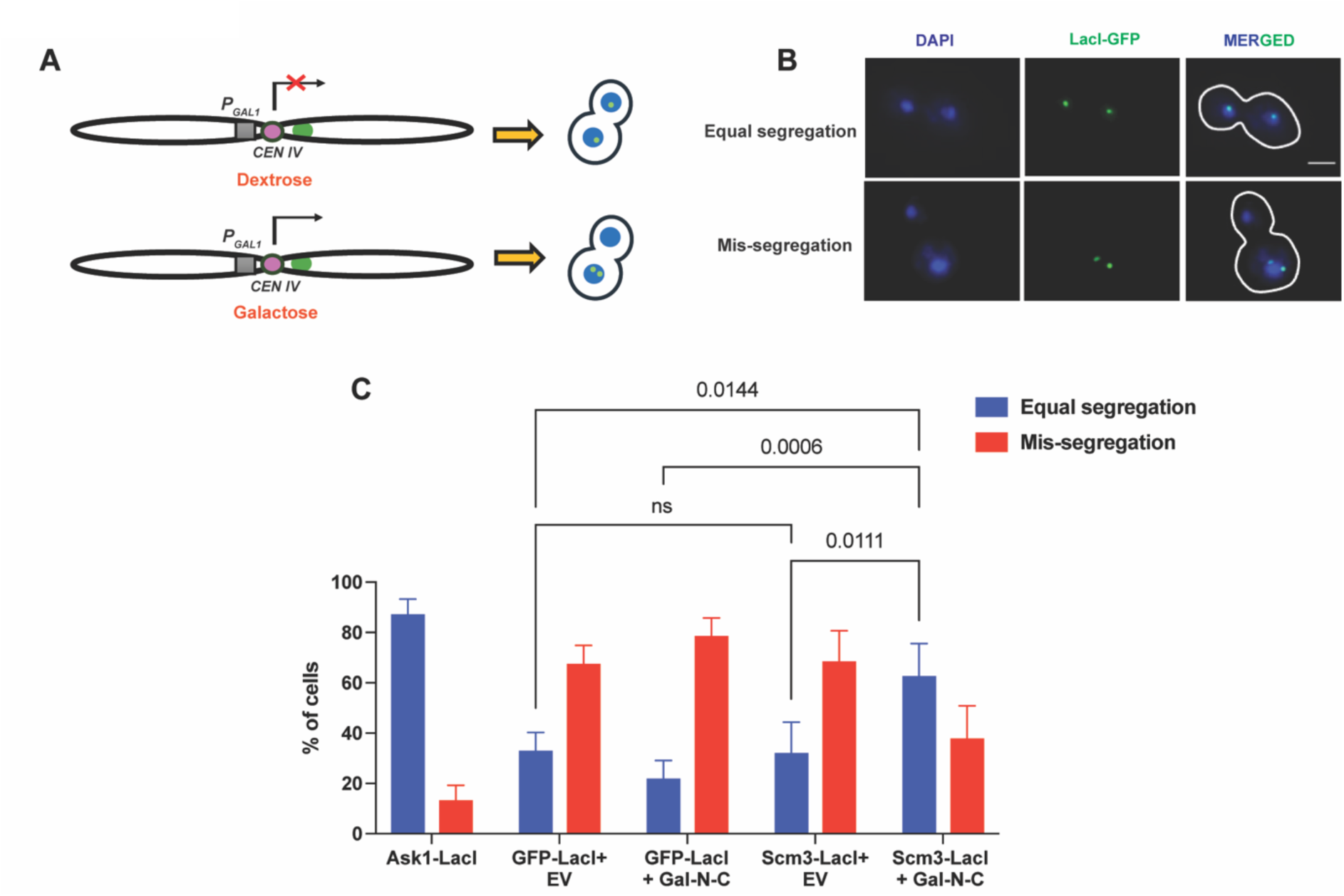
Scm3 along with N-C can drive chromosome segregation. **(A)** Schematic diagram depicting conditional inactivation of *CEN* IV using galactose causing missegregation of chromosome IV. **(B)** Representative images of the anaphase cells showing equal or missegregation of chromosome IV marked by *CEN* IV-GFP. Scale bar = 2 μm **(C)** The percentages of cells used in (B) showing equal segregation or missegregation while expressing the indicated proteins, are presented graphically. N ≥105.

## Discussion

The residence of Scm3 at the centromeres following deposition of Cse4 (Xiao et al., 2011) provided a clue that this protein might have additional functions at the centromeres. In support of this, an interaction between Scm3 and N-C has been reported in recent studies (Shukla et al., 2024, Popchock et al., 2025) where it is proposed that the interaction augments recruitment of AO at the centromeres which has potential to promote kinetochore assembly. In this work, we demonstrate the occurrence of the interaction in vivo and potential of this interaction to rescue equal segregation of a missegregating chromosome possibly by forming active kinetochore.

### The normal stoichiometry of kinetochore proteins is crucial for cell viability

The kinetochore complex of budding yeast is a macromolecular nucleoprotein structure, consisting of more than 100 proteins residing at point centromeres. The complex is subdivided into several sub-complexes, each requiring specific stoichiometry of its component proteins to form a fully functional kinetochore. Misregulation of majority of these proteins often leads to chromosome instability which can result in cell death (Bakhoum and Compton, 2012; Yuen et al., 2005). For example, in budding yeast, the misregulation of Cse4 has been extensively linked to chromosomal instability (CIN) which arises primarily due to misincorporation of Cse4 at the non-centromeric loci (Mishra et al., 2021; Ohkuni et al., 2024; Zhang et al., 2024). Here, we demonstrate that even the N-terminal tail of Cse4 (N-C), when overexpressed, leads to significant loss of cell viability, particularly in kinetochore mutants (Figure 1). Since N-C is known to promote kinetochore assembly through its physical interaction with the essential subunits, Ame1 and Okp1 of the COMA sub-complex (Anedchenko et al., 2019; Fischböck-Halwachs et al., 2019), perhaps these proteins are titrated out by excess N-C and become limiting at the centromeres. The AO limiting condition affected cell viability more in *ctf19Δ* or *mcm21Δ* cells than in *ctf3Δ* or wild type cells because the former cells were already compromised in COMA functions (Figure 1B). Therefore, a typical level of N-C is crucial to maintain the normal stoichiometry of the proteins at the kinetochore supporting cell viability.

On the other hand, overexpression of Scm3 is also implicated in chromosomal instability (Mishra et al., 2011). Excess Scm3 competes with Cse4 for centromeric binding, leading to a reduction in Cse4 occupancy at the centromere and a failure to properly assemble kinetochores (Mishra et al., 2011). Although Scm3 typically interacts with the HFD of Cse4, recent in vitro study has revealed that Scm3 also binds to N-C (Shukla et al., 2024). Further, the interaction between Scm3 and N-C enhances the efficiency of recruitment of the AO to the kinetochore (Shukla et al., 2024). Interestingly, in accord to this, we observed that when both Scm3 and N-C were co-overexpressed, the loss of cell viability was significantly greater than when either of the protein was overexpressed (Figure 2). This was supported by further reduction of centromeric Ame1-13Myc in cells overexpressing both the proteins. The perturbed centromeric localisation of Ame1-13Myc contributes to weaken the kinetochore integrity and loss of cell viability (Figure 3). This synergistic effect strongly suggests a functional interaction between Scm3 and N-C. Thus, we provide first-hand in vivo evidence of this interaction enhancing AO stabilization and thus show how Scm3 might have additional functions beyond Cse4 deposition at the kinetochore.

While this work was in progress, a parallel work, using single molecule fluorescence assay, has reported an in vitro interaction between Scm3 and END domain of N-C which may compensate for the Cse4 maintenance defect arisen due to the loss of END-AO interaction (Popchock et al., 2025). In support of this, we demonstrate the in vivo interaction between Scm3 and N-C using yeast two hybrid and co-immunoprecipitation assays (Figure 4).

### Scm3-N-C interaction also stabilizes Cse4 nucleosome

It is believed that the presence of Scm3 at the centromeres beyond S phase, when Cse4 is deposited, is required to protect Cse4 from random eviction as the removal of Scm3 during anaphase results in the loss of Cse4 from the centromeres (Luconi et al., 2011). However, how Scm3 might provide the protection function is not clear; we believe that the interaction between Scm3 and N-C is crucial in this and perhaps is required to stabilize Cse4 at the centromeres following its deposition. In support of this, the recent study has demonstrated that a ternary interaction among Scm3, N-C and AO stabilizes Cse4 at the centromeres (Popchock et al., 2025).

Intrinsically disordered proteins, such as N-C and Scm3, are capable of forming large macromolecular complexes due to their flexible structures (Shukla et al., 2024). The interaction between these two disordered proteins provides a platform for the deposition of kinetochore subunits, particularly the COMA sub-complex, and enhances the stability of the kinetochore. The dynamic and flexible nature of N-C allows it to span a large area due to its intrinsically disorder nature, which is probably stabilized by the interaction with Scm3 and becomes conducive to subsequent deposition of COMA proteins including AO leading to kinetochore assembly (Figure 6).

**Figure 6:**
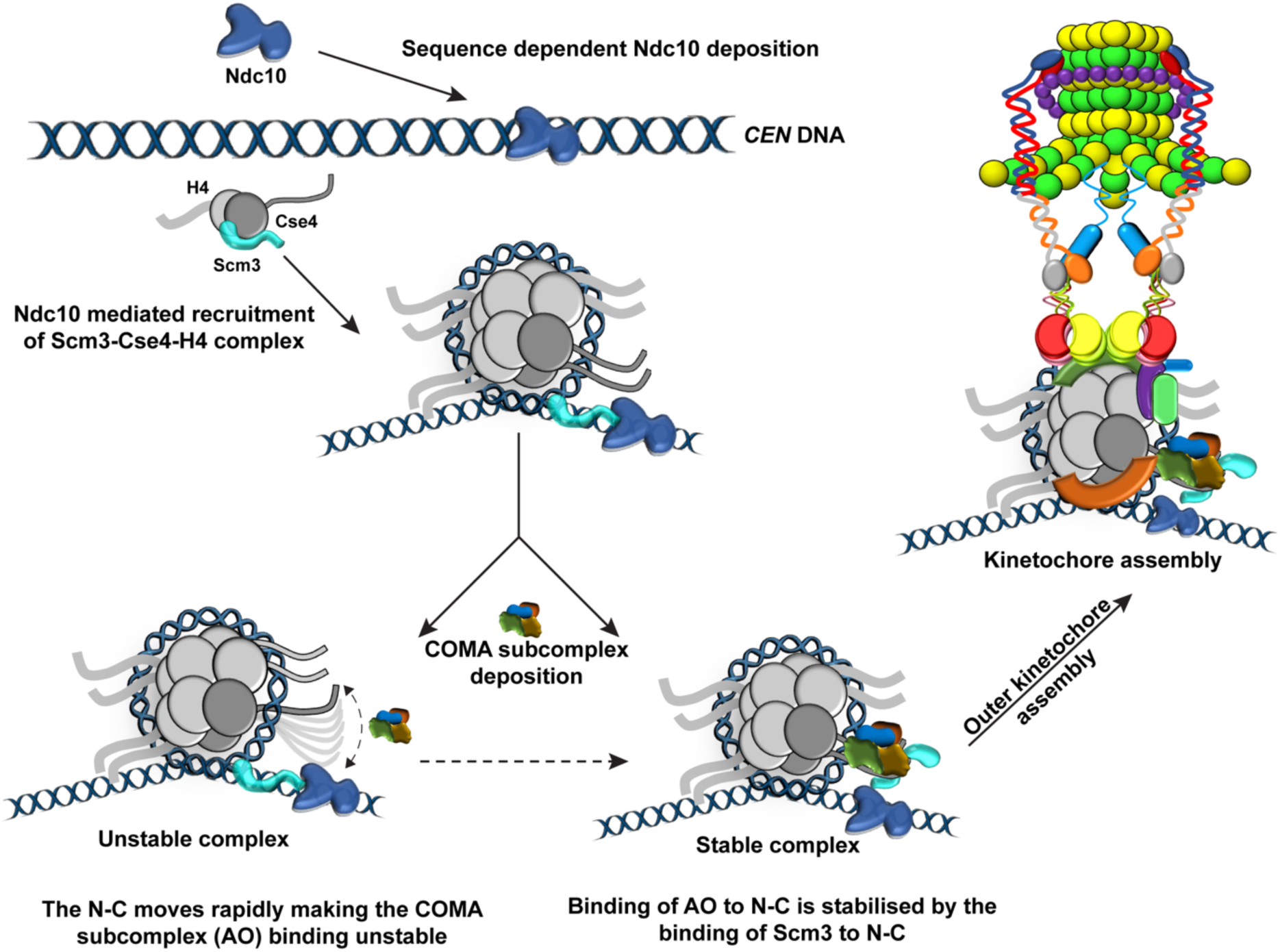
A working model summarising the contribution of the interaction between Scm3 and N-C in kinetochore assembly. The kinetochore assembly is initiated by the sequence-dependent binding of Ndc10 at the centromere. Ndc10 recruits the Cse4-H4-Scm3 complex at the centromere for the formation of centromeric nucleosome. Following Cse4 recruitment at the centromeric nucleosome, the disordered N-C tail moves rapidly making the binding of the COMA subcomplex to N-C difficult creating an unstable state. Scm3 persisting at the centromere interacts with N-C and stabilizes it, facilitating its proper binding to the COMA. This stabilizes the centromeric nucleosome resulting in subsequent assembly of the outer kinetochore complex.

### Scm3-N-C interaction along the cell cycle

It is evident from this and other studies that Scm3 binds in vivo to N-C and helps in the initiation of kinetochore assembly. Thus, the interaction between Scm3 and N-C addresses why and how Scm3 resides at the centromeres even after deposition of Cse4 although the latter can also happen through DNA-binding capacity of Scm3. It is possible that this interaction occurs or peaks up in a cell cycle dependent manner. When kinetochore assembly takes place following deposition of Cse4 at S phase, the interaction initiates perhaps with a peak and then may sustain at a reduced level along the cell cycle maintaining Cse4 at the centromeres. It is possible that the extent of interaction can be modulated at onset of anaphase due to change in microtubule based tension on the kinetochores. Although it is not known whether the same pool of Scm3 that recruits Cse4 at the centromeres, resides throughout the cell cycle, it would be intriguing to test this and find which pool interacts with N-C. Previous in vitro study suggested that N-C becomes available for interaction following Cse4 deposition and formation of Cse4-H4 dimer (Shukla et al., 2024), when Scm3 also dissociates from the HFD of Cse4 as Cse4-H4 is formed. Therefore, in principle, the same Cse4-depositing Scm3 can interact with available N-C. However, given the fast and dynamic exchange of Scm3 in the nucleus (Wisniewski et al., 2014, Luconi et al., 2011), it is quite possible that there is another Scm3 molecule that binds to N-C than the one that deposits Cse4.

### Interaction between Scm3 and N-C : a promoter of kinetochore assembly

We have shown that overexpressed Scm3 and N-C reduce cell viability specifically in kinetochore mutants (Figure 2). We reasoned this phenotype is due to their interaction with key kinetochore proteins sequestering them away from the centromeres. The rescue of equal segregation of a missegregating chromosome upon tethering of Scm3 on it is encouraging to believe that an active kinetochore has been formed at the tethered site (Figure 5). It would be interesting to examine the localization of AO and Cse4 at that site because as per our hypothesis, AO, but not Cse4, should be localized there. The ability of Scm3 and N-C to form an active kinetochore at ectopic sites should be carefully investigated using the homologous proteins in humans as this posits development of aneuploidy particularly when there are reports of genome wide localization of Scm3 in yeast (Prakhar Agarwal and SK Ghosh, unpublished observation) and inadvertent high expression of both HJURP and CENP-A in mammals.

In summary, the misregulation of Scm3 and the N-terminal tail of Cse4 can reduce cell viability by disrupting kinetochore assembly, particularly in cells with pre-existing defects in kinetochore function. The in vivo interaction of Scm3 with N-C is crucial for kinetochore assembly and perhaps required for the maintenance of its stability throughout the cell cycle. Examining this interaction in human cells would be intriguing, where the size of the CENP-A chaperone is much bigger, and multiple CENP-A nucleosomes interspersed with H3-nucleosomes occupy the centromeres. It is possible that the interaction may facilitate to form right 3D organization of the human kinetochores where CENP-A nucleosomes are phased outwards towards the microtubules.

## Data availability

Strains and plasmids used in this work are listed in Supplementary files 1A and 1B, respectively. All data necessary for confirming the conclusions of the article are present within the article, figures, table, and supplementary material.

## Acknowledgement

We thank Wei-Chun Au from National Institutes of Health, USA for providing us with plasmids and YMB11828 strain. This work is supported by Department of Biotechnology (DBT), Govt of India grant (BT/PR43050/BRB/10/1992/2021) to SKG. PA, AA and SM are supported by fellowship grants from CSIR (09/087(0972)/2019-EMR-I), DST (IF230484) and CSIR (09/0087(17614)/2024-EMR-I), Govt of India, respectively. We acknowledge the central instrumentation facility of IIT Bombay for confocal microscopes.

## Materials and Methods

### Strains and plasmids used

All *Saccharomyces cerevisiae* strains used in this study are derivatives of the W303 strain background and are listed in Supplementary file 1A. Specifically, PJ69-4A (James et al., 1996) was utilised for the yeast two-hybrid assay. YMB11828 (Mishra et al., 2023), was utilised for the assessment of kinetochore protein Ame1, in immunofluorescence and ChIP assay. DY6282 (Reid et al., 2008) was modified for the centromere inactivation assay. Plasmids used in the study are listed in Supplementary file 1B. Standard growth conditions were utilized to culture and harvest the yeast cells for all the experiments. To maintain the episomal plasmids, appropriate selection media were used. Gene deletions and C-terminal tagging (AID) were performed using PCR-based gene disruptions or integration at their respective endogenous loci (Janke et al., 2004; Longtine et al., 1998; Wach, 1996). Strains were constructed using the lithium acetate-based yeast transformation method (Gietz and Woods, 2001).

For the yeast-two hybrid assay, pGBD, carrying the *GAL4* DNA binding domain (BD) and pGAD, carrying the *GAL4* activation domain (AD) were utilized. Plasmids with the gene fusions were constructed as follows. The BD-Scm3 (SGB15002) plasmid was constructed by amplifying the coding sequence of *SCM3* from the genome with flanking *Pst*I and *Eco*RI sites. The *Eco*RI*-Pst*I fragment of the PCR product was ligated to pGBD. The AD-Cse4 (SGB15003) plasmid was constructed by amplifying the coding sequence of *CSE4* from the genome with flanking *Pst*I and *Eco*RI sites. The *Eco*RI*-Pst*I fragment of the PCR product was ligated to pGAD. The AD-N-C (SGB15004) plasmid was constructed by amplifying the coding sequence of *CSE4,* with a stop codon after 129 bp from plasmid pMB1234 with flanking *Pst*I and *Eco*RI sites. The *Eco*RI*-Pst*I fragment of the PCR product was ligated to pGAD.

For galactose inducible overexpression of N-C, *Pvu*II fragment containing *N-C* from pMB1234 was ligated with the *Pvu*II fragment containing *URA3* marker from pMB1236, to construct SGB10112. For galactose inducible overexpression of Myc tagged N-C, *Pvu*II fragment containing *N-C-MYC* from pMB1236 was ligated with the *Pvu*II fragment containing *LEU2* marker from pMB1193, to construct SGB15001.

To construct the LacO-LacI based artificial kinetochore system within *S.cerevisiae* the strain construction was done as follows (Lacefield et al., 2009; Murray and Szostak, 1983). The pRS-403-GFP-LacI plasmid (6.4 kbp) was modified with Klenow treatment at the *Xho*I site flanking LacI gene followed by sequential digestion with *Eco*RI and *Xho*I, the larger fragment of 1.129 kbp was gel purified for vector backbone while the *GFP* gene containing smaller fragment is excluded. The coding sequence of *SCM3* gene was amplified from the genome with flanking sites for *Eco*RI and *Xho*I. The *Eco*RI*-Xho*I containing PCR product of *SCM3* is ligated onto 1.129 kbp fragment of the modified pRS-403-GFP-LacI plasmid to construct SGB14003 (∼6.5 kbp) with *SCM3-LacI* fusion construct (Kiermaier et al., 2009; Lacefield et al., 2009).

### Media, reagents, and growth conditions

Yeast strains were cultured using standard growth conditions (Sherman, 2002). Appropriate selection media were used to maintain the episomal plasmids. For galactose inducible overexpression, strains were cultured in 2% raffinose containing selection medium prior to induction with 2% galactose. For plates containing 3-AT (Himedia, India) and Auxin (Sigma Chemicals, Switzerland), the compounds were surface spread onto selection plates at indicated concentration. To arrest the cells at metaphase, nocodazole (Sigma Chemicals, Switzerland) was added to the cultures at indicated concentrations. For western blotting, immunoprecipitation and immunofluorescence studies, rat anti-HA (3F10, Roche), mouse anti-HA (12CA5, Roche), mouse anti-Myc (9E10, Roche), rat anti-Tubulin (MCA78G, Serotec), were utilized as the primary antibodies, at appropriate dilutions. Rhodamine (TRITC)-conjugated goat anti-Rat (Jackson), Alexa Fluor 488 conjugated goat anti-Mouse IgG (Jackson), HRP-conjugated goat anti-Mouse IgG (Jackson), and HRP-conjugated goat anti-Rat IgG (Jackson), were utilized as the secondary antibodies at appropriate dilutions.

### Cell spotting assay and cell viability assay

Strains were grown to mid-log phase in the indicated selection media containing 2% raffinose. The cultures were spotted in 10-fold serial dilutions on indicated selection plates containing either 2% galactose or 2% dextrose. All indicated reagents were surface plated on the agar plates. Plates were incubated at 30°C for 2 days or at 16°C or 11°C for 2-5 days, according to the growth rate and were photographed.

For measuring the cell viability, strains were grown to mid-log phase in indicated selection media containing 2% raffinose. Ten-fold serial dilutions were made, and cells were counted using a haemocytometer, and equal number of cells were plated onto appropriate selective plates containing either 2% dextrose or 2% galactose. The plates were incubated at 30°C for 2 days and at 16°C or 11°C 1 for 2–5 days, according to the growth rate. The percentage cell viability was measured as the number of colonies that showed visible growth after the indicated incubation period. Colonies formed on the dextrose plate was treated as 100% cell viability. To test the functionality of *SCM3* constructs (*Scm3-LacI and BD-SCM3*) and *CSE4* construct (*AD-CSE4),* AID tagged strains were serially diluted and spotted onto plates containing either 0.5mM auxin or DMSO. The plates were incubated at 30°C for 2–3 days and were photographed.

### Yeast-two hybrid assay and β-galactosidase assay

Plasmids pGAD and pGBD, with or without the respective gene inserts (*N-C*, *CSE4*, *SCM3*), were co-transformed into the strain PJ69-4A, in different combinations and plated onto synthetic complete plates lacking leucine and tryptophan. The transformants were streaked onto synthetic complete plates lacking leucine, tryptophan, and histidine, with or without different concentrations of 3-AT. To quantitatively measure the interaction, β-galactosidase activity was measured using ONPG (o-nitrophenyl-β-D-galactopyranoside, ThermoFisher) as a substrate. The activity was calculated in Miller units.

### Co-immunoprecipitation

Overnight grown culture in synthetic complete medium lacking leucine and uracil with 2% raffinose was diluted to 0.4 OD in 200 mL of synthetic complete medium lacking leucine and uracil supplemented with either 2% galactose or 2% dextrose. The cells were grown till 0.8 OD. The cells were harvested by centrifugation at 3000 rpm for 2 minutes at 4°C. The pellet was washed with 30 mL of ice-cold water. The pellet was washed once with lysis buffer supplemented with 1X PIC, 1 mM PMSF. The cell pellet was resuspended in 1 mL of lysis buffer supplemented with 1X PIC and 1 mM PMS, and distributed into 2 O-ring tubes (around ∼600 µL each). The cells were lysed with 0.5 mm glass beads using a mini-bead beater (BIOSPEC, India) for 7 cycles (1 min ON, 2 mins OFF on ice). After ∼80% of cell lysis was observed, the O-ring tube was punctured and kept in an inverted position in a 15 mL falcon. The total lysate was collected after spinning the falcons at 2800 rpm for 2 minutes at 4°C. The lysate was clarified by spinning at 13,500 rpm for 15 minutes at 4°C. The supernatant was collected and an input fraction was assessed by normal western blotting for the presence of epitope tagged protein. The rest was split into two portions, one with the ∼ 5 µg antibody (ab9110, ab9106, Abcam) and the other without any antibody. The tubes were incubated at 4°C, overnight, with rotation. The lysates were then incubated with 12 µL of Dynabeads Protein G (pre-equilibrated in the lysis buffer), for 4 hours with rotation. After collecting the beads by placing on a magnetic stand for 2 minutes, the supernatant was collected as the unbound fraction. To the beads, lysis buffer was added and rotated for 5 minutes, The supernatant collected after placing on the magnetic stand, the tubes was stored as wash I. After three such washes, the beads were resuspended SDS buffer and boiled at 95°C for. The eluate was collected by placing the tubes on a magnetic stand and separating the eluate from the beads.

### Centromere inactivation assay

Strain DY6282 with conditionally inactivated *CEN IV* (Reid et al., 2008), was used for the centromere inactivation assay. To mark centromere IV, 256 LacO repeats were added at the *TRP1* locus and GFP-LacI-LacO system was used to track the segregation of *CEN IV* and the strain was transformed with Scm3-LacI, with or without overexpression of N-C-Myc. Strain transformed with Ask1-LacI (Lacefield et al., 2009) construct was used as a positive control. An overnight grown culture in synthetic media lacking leucine, supplemented with 2% raffinose was diluted to an OD of 0.2 in synthetic media lacking leucine, supplemented with 2% galactose, and allowed to grow till 0.8-1 OD. Cells were harvested and treated with DAPI to image the cells and study the segregation of GFP marked *CEN IV* with respect to the segregation of DAPI stained nuclei.

### Western blotting

Standard procedures were followed for performing western blotting. Ame1-Myc and N-C-Myc were detected using mouse anti-Myc antibody (9E10, Roche) and Scm3-HA was detected using rat anti-HA antibody (3F10, Roche). HRP-conjugated goat anti-Mouse IgG (Jackson), and HRP-conjugated goat anti-Rat IgG (Jackson), were used as the secondary antibodies at appropriate dilutions.

### Indirect immunofluorescence

Immunofluorescence was performed as mentioned previously (Anbalagan et al., 2024) Briefly, culture was harvested after nocodazole mediated metaphase arrest and fixed with 4% formaldehyde for 15 minutes at room-temperature (RT). After fixation, the cells were washed twice with 0.1M Phosphate buffer and spheroplasted with spheroplasting solution (1.2M sorbitol, 0.1M phosphate buffer at pH 7.5, Zymolase 20T; 10mg/mL, 25mM β-Mercaptoethanol) for 30 minutes at 30°C. The spheroplasts were washed with spheroplasting buffer and PBS and resuspended in PBS. The spheroplasts were mounted onto poly-L-lysine coated slide and incubated at RT for 30 minutes. The spheroplasts were flattened by immersing the slides in pre-chilled (-20°C) methanol and acetone for 5 minutes and 30 seconds, respectively. Slides were blocked using 5% skimmed milk in 10 mg/mL BSA in PBS, for 1 hour. After washes with PBS, the spheroplasts were incubated with the primary antibody at 1:300 dilution and secondary antibody at 1:400 dilution, for 1 hour each at RT. Following PBS washes, the spheroplasts were incubated with DAPI (Invitrogen, #D1306) (1 μg/ml) solution for 15 minutes in the dark. The slides were layed with mounting medium (1 mg/mL phenylenediamine in 90% glycerol) and sealed with a coverslip.

### Fluorescence microscopy

All fluorescence images were captured using the Zeiss Axio Observer.Z1 (100X objective, NA=1.40; Carl Zeiss Micro-Imaging Inc.) equipped with Axiocam camera. Further, the images were processed and analysed using the Zeiss Zen 3.1 software.

### ChIP (Chromatin immunoprecipitation) assay and qPCR quantification

Chromatin immunoprecipitation was performed as previously mentioned (Anbalagan et al., 2024, Kumar et al., 2021). First, cells were grown in the presence of 2% raffinose, in appropriate selective media and then transferred to selection media containing 2% galactose. At 0.6 OD, nocodazole was added to arrest cells at metaphase. A total of 60 OD cells were harvested The cells were then fixed with 1% formaldehyde for 30 mins at 25°C at 100 rpm, and then quenching was performed by adding glycine at a final concentration of 125 mM. The cells were then harvested and washed twice in ice-cold TBS buffer (20 mM Tris–HCl pH 7.5, 150 mM NaCl) and once with 10 ml of FA lysis buffer (50 mM Hepes–KOH pH 7.5, 150 mM NaCl, 1 mM EDTA, 1% Triton X-100, 0.1% Na-deoxycholate) containing 0.1 % SDS. The pellet was resuspended in 500 µl of 1X FA lysis buffer supplemented with 0.5% SDS, 1 mM PMSF, 1X PIC. The cells were lysed using 0.5 mm glass beads using a mini-bead beater (BIOSPEC, India) for 7 cycles (1 min ON, 2 mins OFF on ice). The lysate was collected in a pre-chilled 1.5 ml eppendorf tube and centrifuged twice at 16,000 x g for 15 mins at 4°C. The pellet was resuspended in 300 µl ice-cold 1X FA lysis buffer supplemented with 0.1% SDS, 1 mM PMSF, and 1X PIC. The chromatin was then sheared to 200-600 bps using a water bath sonicator (Diagenode SA, Picoruptor, BC 100,27 LAUDA Germany) for 30 cycles of 30 s ON / 30 s OFF at 4°C. After clarification of the lysate by centrifugation at 16,000 x g for 10 mins at 4°C, 3-5 µg of appropriate antibodies (were added and incubated at 4°C overnight with gentle rotation. Subsequently, 50 µl Protein-A conjugated Sepharose beads were added and incubated for 2 hours at 4°C with rotation. The beads were washed twice with IP wash buffer I (1X FA lysis buffer, 0.1%SDS, 275 mM NaCl), followed by one wash in IP wash buffer II (1X FA lysis buffer, 0.1%SDS, 500 mM NaCl, III (10 mM Tris pH 8.0, 250 mM LiCl, 0.5% NP-40, 0.5% sodium deoxycholate, 1 mM EDTA), and 1X TE at room temperature The beads were resuspended in elution buffer and the chromatin was eluted by boiling at 65°C. The eluate was then de-crosslinked overnight at 65°C, followed by Proteinase K (SRL, India) treatment for 2 hours at 45°C. The DNA was purified by PCI-based purification and precipitated overnight in chilled ethanol at -80°C. The enrichment of obtained chromatin fragments was estimated using qPCR (BioRad CFX96, USA) using specific primers targeting specific and non-specific (negative control) chromatin loci, listed in Supplementary file 1C. As mentioned before (Kumar et al., 2021; Shah et al., 2023) the following equation was used to estimate the percentage of chromatin enrichment. ChIP efficiency = Enrichment/Input X 100; Enrichment/Input = E˄-ΔCT; ΔCT = CT(ChIP) − [CT(Input) − LogEX(D)]; E = primer efficiency value, CT = Threshold values obtained from qPCR, D = Input dilution factor. E was estimated as {[10^(-1/slope)]– 1} from standardization graphs of CT values against dilutions of the input DNA.

### Statistical analysis

The data presented here were obtained from two to three independent experiments. The error bars in the individual bar graphs represent the standard error (SE) derived from the standard deviation (SD). The statistical significance (*p*) was determined by two-tailed Student’s t-test, or one-way ANOVA test, as appropriate. When comparing more than one group, a one-way ANOVA test was used; otherwise, Student’s t-test was used. The ‘N’ values denote the total number of cells analysed from the combined ‘n’ number of replicates of the individual assays. The *p* values less than or equal to 0.05 are categorized as significant differences. The SD, SE, and statistical significance (*p*) values were calculated using automated modules of GraphPad Prism 9.0 (Version 9.4.1) software.

---

**Figure S1:**
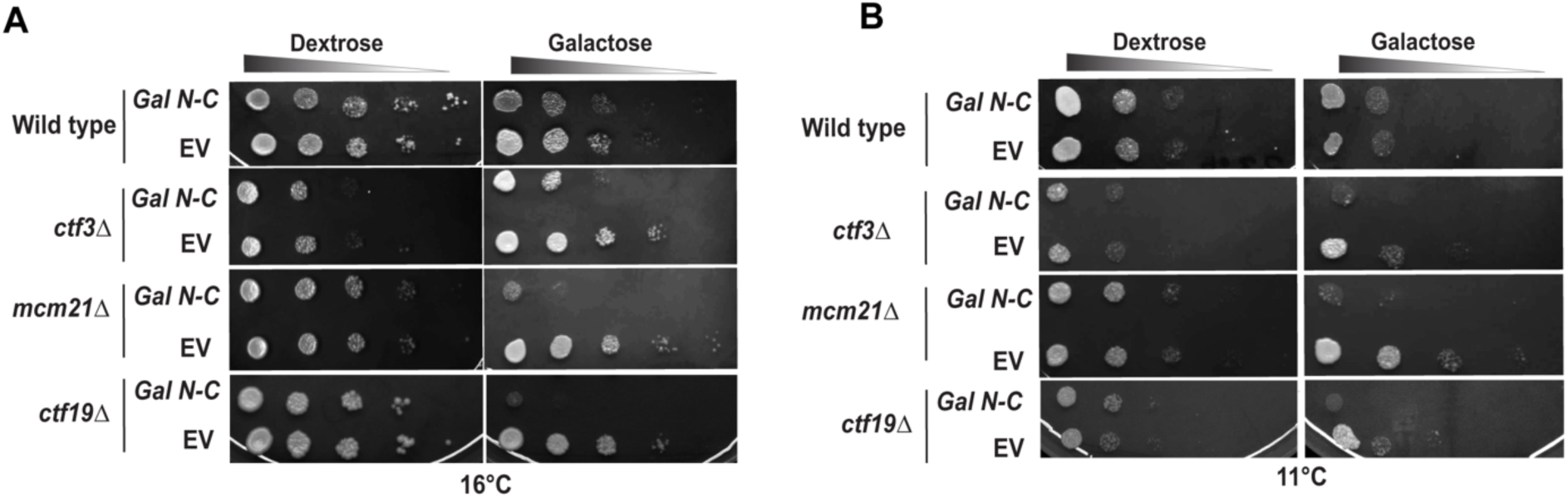
Overexpression of N-C leads to loss of cell viability in kinetochore mutants at lower temperatures. Wild-type, *ctf3Δ, mcm21Δ, and ctf19Δ* cells harboring *Gal N-C* or empty vector (EV) were grown till mid-log before they were serially diluted and spotted on the indicated plates. The plates were incubated at 16°C **(A)** or 11°C **(B)** for 48-60 hrs before imaging.

**Figure S2:**
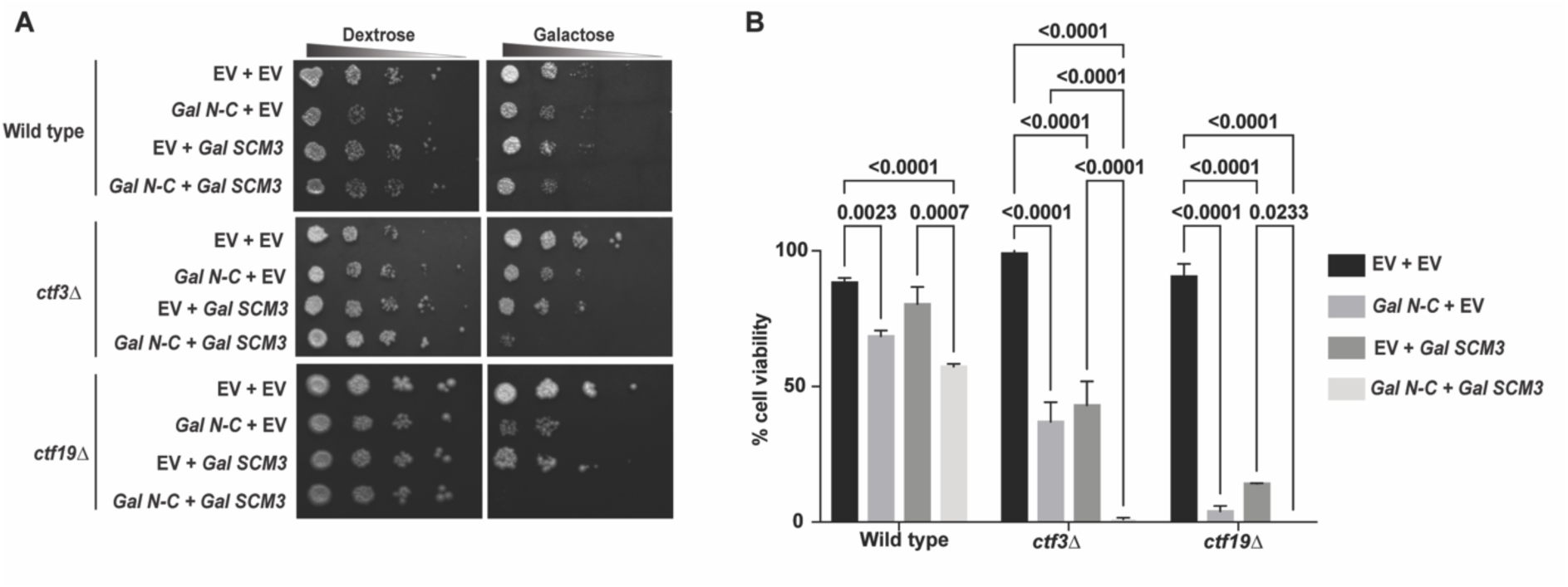
Overexpression of both N-C and Scm3 causes synthetic sick phenotype at lower temperature. Same as Figure 2, except the plates were incubated at 16°C.

**Figure S3:**
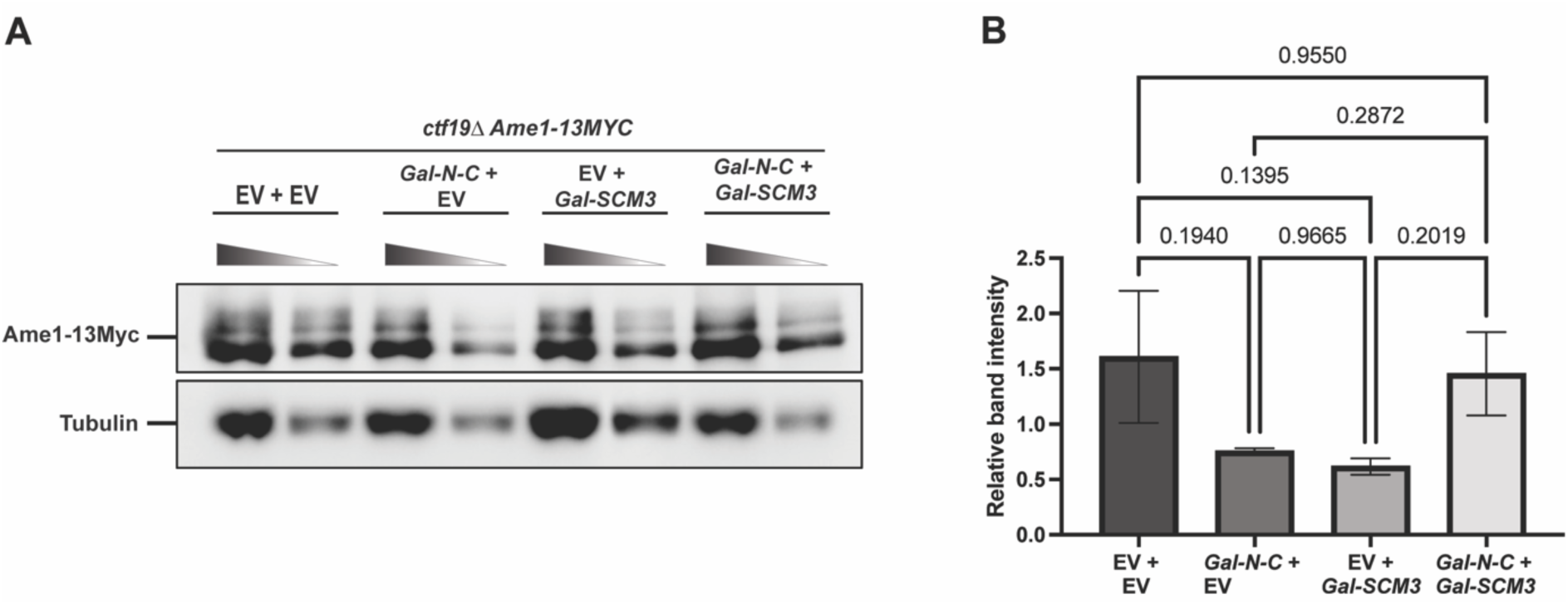
The overexpression of N-C and Scm3 does not affect Ame1 expression. **(A)** Western blots show the levels (in 2-fold serial dilutions) of Ame1-13Myc in metaphase arrested cells used in Figure 3A. Anti-Myc and anti-tubulin antibodies were used to detect the corresponding proteins. **(B)** The graph shows the relative band intensity measured from the average of the two band intensities for each sample normalized with the background and average of the corresponding two tubulin bands. Error bars represent the standard deviation from the mean values obtained from two independent experiments. The statistical significance p value was calculated using a two-way ANOVA test.

**Figure S4:**
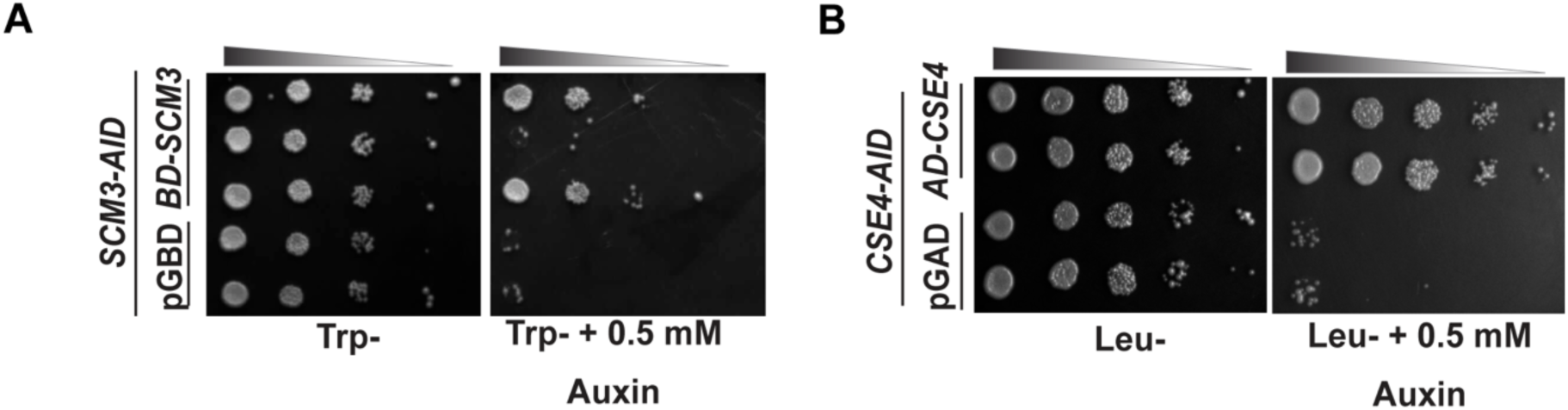
BD-Scm3 and AD-Cse4 fusion proteins are functional. **(A)** Yeast strain harboring *SCM3-AID* as the sole source of *SCM3* was transformed with either BD-*SCM3* fusion construct or empty vector (pGBD). Three independent transformants were spotted on the indicated plates in the presence or absence of auxin to degrade Scm3. The plates were incubated at 30°C for 3-4 days before imaging. **(B)** Same as (A), except strain harboring *CSE4-AID* as the sole source of *CSE4* was transformed by AD-*CSE4* fusion construct or empty vector (pGAD), and two independent transformants were spotted.

**Figure S5:**
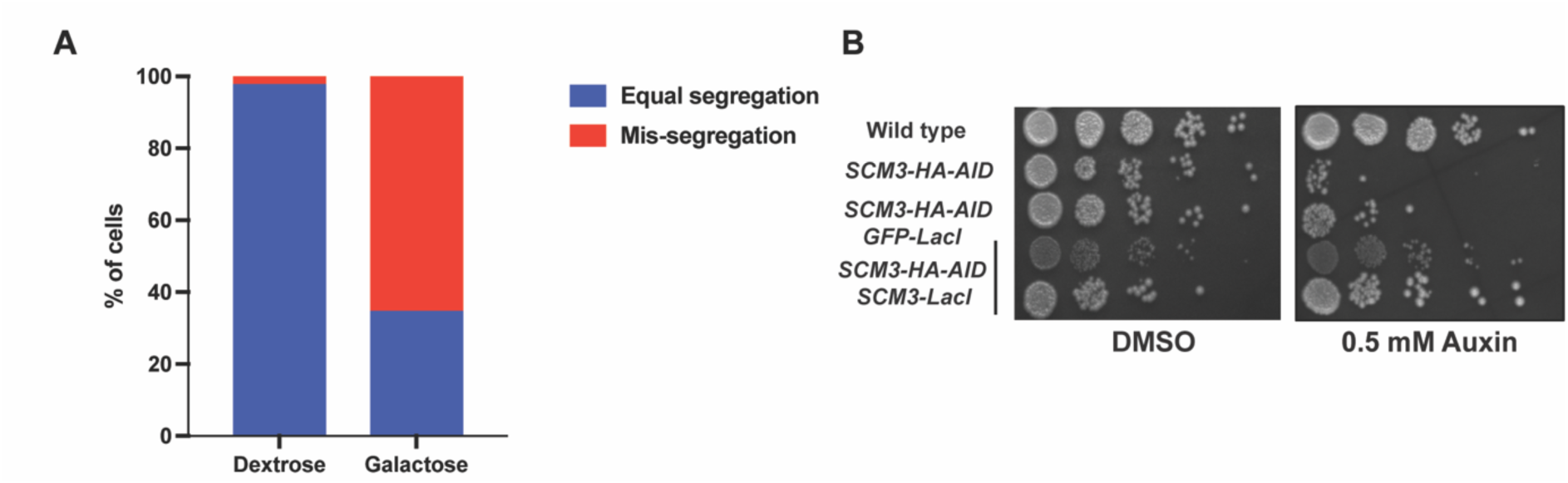
Validation of conditional inactivation of centromere in galactose and functionality of Scm3-LacI fusion protein. **(A)** The percentages of cells showing equal segregation or missegregation of chromosome IV grown under dextrose or galactose are shown graphically. N = 98. **(B)** The yeast strains with indicated genotypes were serially diluted (1:10) and spotted on YPD palates with or without (DMSO) auxin. The growth of the Scm3 depleted cells (auxin panel) can be rescued by Scm3-LacI (4^th^ and 5th rows) but not by GFP-LacI (3^rd^ row).

## Supplementary material

**Supplementary file 1A: List of strains used in this study:**

| S.No. | Strain code | Genotype (all strains are of W303 background) | Reference |
| --- | --- | --- | --- |
| 1 | CRY1 | *MATa, ura3-1 leu2,3-112 his3-1 trp1-1 ade2-1 can1-100* | Rothstein, 1983 |
| 2 | SGY10137 | *MATa ade2-1 trp1-1 can1-100, leu2-3, 112 his3-11,15 ura3 pGAL1-N-C-13MYC::URA3 (PMB1236)* | This study |
| 3 | SGY10528 | *MATa ade2-1 trp1-1 can1-100, leu2-3, 112 his3-11,15 ura3 pESC-URA3::URA3* | This study |
| 4 | SGY10124 | *MATa ade2-1 trp1-1 can1-100, leu2-3, 112 his3-11,15 ura3, ctf3Δ::HPHMX pGAL1-N-C-13MYC::URA3 (PMB1236)* | This study |
| 5 | SGY10125 | *MATa ade2-1 trp1-1 can1-100, leu2-3, 112 his3-11,15 ura3, ctf3Δ::HPHMX pESC-URA3::URA3* | This study |
| 6 | SGY10156 | *MATa ade2-1 trp1-1 can1-100, leu2-3, 112 his3-11,15 ura3, ctf19Δ::HPHMX pGAL1-N-C-13MYC::URA3 (PMB1236)* | This study |
| 7 | SGY10155 | *MATa ade2-1 trp1-1 can1-100, leu2-3, 112 his3-11,15 ura3, ctf19Δ::HPHMX pESC-URA3::URA3* | This study |
| 8 | SGY10126 | *MATa ade2-1 trp1-1 can1-100, leu2-3, 112 his3-11,15 ura3, mcm21Δ::HPHMX pGAL1-N-C-13MYC::URA3 (PMB1236)* | This study |
| 9 | SGY10127 | *MATa ade2-1 trp1-1 can1-100, leu2-3, 112 his3-11,15 ura3, mcm21Δ::HPHMX pESC-URA3::URA3* | This study |
| 10 | SGY10139 | *MATa ade2-1 trp1-1 can1-100, leu2-3, 112 his3-11,15 ura3 pESC-URA3::URA3 pESC-LEU2::LEU2* | This study |
| 11 | SGY10140 | *MATa ade2-1 trp1-1 can1-100, leu2-3, 112 his3-11,15 ura3 pESC-URA3::URA3 pGAL1-SCM3 (PMB1193)* | This study |
| 12 | SGY10141 | *MATa ade2-1 trp1-1 can1-100, leu2-3, 112 his3-11,15 ura3 pGAL1-N-C-13MYC::URA3 (PMB1236) pESC-LEU2::LEU2* | This study |
| 13 | SGY10142 | *MATa ade2-1 trp1-1 can1-100, leu2-3, 112 his3-11,15 ura3 pGAL1-N-C-13MYC::URA3 (PMB1236) pGAL1-SCM3 (PMB1193)* | This study |
| 14 | SGY10186 | *MATa ade2-1 trp1-1 can1-100, leu2-3, 112 his3-11,15 ura3, ctf19Δ::HPHMX pESC-URA3::URA3 pESC-LEU2::LEU2* | This study |
| 15 | SGY10185 | *MATa ade2-1 trp1-1 can1-100, leu2-3, 112 his3-11,15 ura3, ctf19Δ::HPHMX pESC-URA3::URA3 pGAL1-SCM3 (PMB1193)* | This study |
| 16 | SGY10183 | *MATa ade2-1 trp1-1 can1-100, leu2-3, 112 his3-11,15 ura3, ctf19Δ::HPHMX pGAL1-N-C-13MYC::URA3 (PMB1236) pESC-LEU2::LEU2* | This study |
| 17 | SGY10184 | *MATa ade2-1 trp1-1 can1-100, leu2-3, 112 his3-11,15 ura3, ctf19Δ::HPHMX pGAL1-N-C-13MYC::URA3 (PMB1236) pGAL1-SCM3 (PMB1193)* | This study |
| 18 | SGY10145 | *MATa ade2-1 trp1-1 can1-100, leu2-3, 112 his3-11,15 ura3, ctf3Δ::HPHMX pESC-URA3::URA3 pESC-LEU2::LEU2* | This study |
| 19 | SGY10146 | *MATa ade2-1 trp1-1 can1-100, leu2-3, 112 his3-11,15 ura3, ctf3Δ::HPHMX pESC-URA3::URA3 pGAL1-SCM3 (PMB1193)* | This study |
| 20 | SGY10143 | *MATa ade2-1 trp1-1 can1-100, leu2-3, 112 his3-11,15 ura3, ctf3Δ::HPHMX pGAL1-N-C-13MYC::URA3 (PMB1236) pESC-LEU2::LEU2* | This study |
| 21 | SGY10144 | *MATa ade2-1 trp1-1 can1-100, leu2-3, 112 his3-11,15 ura3, ctf3Δ::HPHMX pGAL1-N-C-13MYC::URA3 (PMB1236) pGAL1-SCM3 (PMB1193)* | This study |
| 22 | YMB11828 | *MATa ura3Δ0 leu2Δ0 his3Δ1 met15Δ0 6His-3HA-CSE4::mx4NAT Ame1-Myc::TRP1* | Mishra et al., 2023 |
| 23 | SGY10199 | *MATa ura3Δ0 leu2Δ0 his3Δ1 met15Δ0 6His-3HA-CSE4::mx4NAT Ame1-Myc::TRP1 ctf19Δ::HPHMX pGAL1-N-C::URA3 pGAL1-SCM3 (PMB1193)* | This study |
| 24 | SGY10197 | *MATa ura3Δ0 leu2Δ0 his3Δ1 met15Δ0 6His-3HA-CSE4::mx4NAT Ame1-Myc::TRP1 ctf19Δ::HPHMX pESC-URA3::URA3 pESC-LEU2::LEU2* | This study |
| 25 | SGY18050 | *MATa ura3Δ0 leu2Δ0 his3Δ1 met15Δ0 6His-3HA-CSE4::mx4NAT Ame1-Myc::TRP1 ctf19Δ::HPHMX pGAL1-N-C::URA3 pESC-LEU2::LEU2* | This study |
| 26 | SGY18051 | *MATa ura3Δ0 leu2Δ0 his3Δ1 met15Δ0 6His-3HA-CSE4::mx4NAT Ame1-Myc::TRP1 ctf19Δ::HPHMX pESC-URA3::URA3 pGAL1-SCM3 (PMB1193)* | This study |
| 27 | SGY18052 | *MATa ade2-1 trp1-1 can1-100, leu2-3, 112 his3-11,15 SCM3-HA-AID::KanMX pADH1-OsTIR1::URA3 pGBD::TRP1* | This study |
| 28 | SGY18053 | *MATa ade2-1 trp1-1 can1-100, leu2-3, 112 his3-11,15 SCM3-HA-AID::KanMX pADH1-OsTIR1::URA3 pGBD-SCM3::TRP1* | This study |
| 29 | SGY18054 | *MATa ade2-1 trp1-1 can1-100, leu2-3, 112 his3-11,15 CSE4-HA-AID::KanMX pADH1-OsTIR1::URA3 pGAD::LEU2* | This study |
| 30 | SGY18055 | *MATa ade2-1 trp1-1 can1-100, leu2-3, 112 his3-11,15 CSE4-HA-AID::KanMX pADH1-OsTIR1::URA3 pGAD-CSE4::LEU2* | This study |
| 31 | pJ694A | *MATa trp1-901 leu2,3-112 ura3-52 his3Δ200 gal4Δ gal80Δ LYS2::GAL1-HIS3 GAL2-ADE2 met2::GAL7-lacZ* | James et al., 1996 |
| 32 | SGY18056 | *MATa trp1-901 leu2,3-112 ura3-52 his3Δ200 gal4Δ gal80Δ LYS2::GAL1-HIS3 GAL2-ADE2 met2::GAL7-lacZ pGAD::LEU2 pGBD::TRP1* | This study |
| 33 | SGY18057 | *MATa trp1-901 leu2,3-112 ura3-52 his3Δ200 gal4Δ gal80Δ LYS2::GAL1-HIS3 GAL2-ADE2 met2::GAL7-lacZ pGAD::LEU2 pGBD-SCM3::TRP1* | This study |
| 34 | SGY18058 | *MATa trp1-901 leu2,3-112 ura3-52 his3Δ200 gal4Δ gal80Δ LYS2::GAL1-HIS3 GAL2-ADE2 met2::GAL7-lacZ pGAD-CSE4::LEU2 pGBD::TRP1* | This study |
| 35 | SGY18059 | *MATa trp1-901 leu2,3-112 ura3-52 his3Δ200 gal4Δ gal80Δ LYS2::GAL1-HIS3 GAL2-ADE2 met2::GAL7-lacZ pGAD-N-C::LEU2 pGBD::TRP1* | This study |
| 36 | SGY18060 | *MATa trp1-901 leu2,3-112 ura3-52 his3Δ200 gal4Δ gal80Δ LYS2::GAL1-HIS3 GAL2-ADE2 met2::GAL7-lacZ pGAD-CSE4::LEU2 pGBD-SCM3::TRP1* | This study |
| 37 | SGY18061 | *MATa trp1-901 leu2,3-112 ura3-52 his3Δ200 gal4Δ gal80Δ LYS2::GAL1-HIS3 GAL2-ADE2 met2::GAL7-lacZ pGAD-N-C::LEU2 pGBD-SCM3::TRP1* | This study |
| 38 | SGY10200 | *MATa ade2-1 trp1-1 can1-100, leu2-3, 112 his3-11,15 ura3 pGAL1-SCM3-3HA::URA3 (PMB1306) pGAL1-N-C-13MYC::LEU2* | This study |
| 39 | DY6282 | *MATa CEN4::Gal1-CEN4-URA3 ade2-1 can1-100 his3-11,15 leu2-3,112 lys2 met17 trp1-1 ura3-1 rad5-535* | Reid et al., 2008 |
| 40 | SGY10540 | *MATa, ura3-1 leu2,3-112 his3-1 trp1-1 ade2-1 can1-100 SCM3-HA-AID::KanMx, pADH-Os-TIR1::URA3* | This study |
| 41 | SGY10558 | *MATa, ura3-1 leu2,3-112 his3-1 trp1-1 ade2-1 can1-100 SCM3-HA-AID::KanMx, pADH-Os-TIR1::URA3 pHIS3-SCM3-LacI::HIS3* | This study |
| 42 | SGY16011 | *MATa, ura3-1 leu2,3-112 his3-1 trp1-1 ade2-1 can1-100 SCM3-HA-AID::KanMx, pADH-Os-TIR1::URA3 pHIS3-GFP-LacI::HIS3* | This study |
| 43 | SGY18035 | *MATa CEN4::Gal1-CEN4-URA3 ade2-1 can1-100 his3-11,15 leu2-3,112 lys2 met17 trp1-1 ura3-1 rad5-535 GFP-LacI::ADE2 LacO::TRP1* | This study |
| 44 | SGY18063 | *MATa CEN4::Gal1-CEN4-URA3 ade2-1 can1-100 his3-11,15 leu2-3,112 lys2 met17 trp1-1 ura3-1 rad5-535 GFP-LacI::ADE2 LacO::TRP1 SCM3-LacI::HIS3 pGAL1-N-C-13MYC::LEU2* | This study |
| 45 | SGY18064 | *MATa CEN4::Gal1-CEN4-URA3 ade2-1 can1-100 his3-11,15 leu2-3,112 lys2 met17 trp1-1 ura3-1 rad5-535 GFP-LacI::ADE2 LacO::TRP1 SCM3-LacI::HIS3 pESC-LEU2::LEU2* | This study |
| 46 | SGY18065 | *MATa CEN4::Gal1-CEN4-URA3 ade2-1 can1-100 his3-11,15 leu2-3,112 lys2 met17 trp1-1 ura3-1 rad5-535 GFP-LacI::ADE2 LacO::TRP1 pGAL1-N-C-13MYC::LEU2* | This study |
| 47 | SGY18066 | *MATa CEN4::Gal1-CEN4-URA3 ade2-1 can1-100 his3-11,15 leu2-3,112 lys2 met17 trp1-1 ura3-1 rad5-535 GFP-LacI::ADE2 LacO::TRP1 pESC-LEU2::LEU2* | This study |
| 48 | SGY18067 | *MATa CEN4::Gal1-CEN4-URA3 ade2-1 can1-100 his3-11,15 leu2-3,112 lys2 met17 trp1-1 ura3-1 rad5-535 GFP-LacI::ADE2 LacO::TRP1 ASK1-LacI::HIS3* | This study |

**Supplementary file 1B: List of plasmids used in this study:**

| Plasmids | Salient features | Source / References |
| --- | --- | --- |
| pESC-URA3 | Yeast episomal vector with a LEU2 marker | Stratagene |
| pESC-LEU2 | Yeast episomal vector with a URA3 marker | Stratagene |
| pMB1236 (URA3) | 2 μm, GAL1/10-N-C-13Myc | Munira A. Basrai |
| pMB1193 (LEU2) | 2 μm, GAL1/10-SCM3 | Mishra et al.,2011 |
| pMB1234 (TRP1) | 2 μm, GAL1/10-N-C | Munira A. Basrai |
| pMB1306<br>(*URA3*) | 2 μm, *GAL1/10-SCM3-3HA* | Mishra et al., 2011 |
| SGB10112<br>(*URA3*) | 2 μm, *GAL1/10-N-C* | This study |
| SGB15001<br>(*LEU2*) | 2 μm, *GAL1/10-N-C-13Myc* | This study |
| pGBD (*TRP1*) | *GAL4* DNA binding domain (BD) | James et al., 1996 |
| pGAD (*LEU2*) | *GAL4* activation domain (AD) | James et al., 1996 |
| SGB15004<br>(*LEU2*) | pGAD-N-C; N-C cloned in pGAD vector for two-hybrid assay | This study |
| SGB15003<br>(*LEU2*) | pGAD-CSE4; CSE4 cloned in pGAD vector for two-hybrid assay | This study |
| SGB15002<br>(*TRP1*) | pGBD-SCM3; SCM3 cloned in pGBD vector for two-hybrid assay | This study |
| pASF52 (*TRP1*) | LacO TRP | Straight et al., 1996 |
| pRS402 GFP-LacI (*ADE2*) | pRS402-GFP-LacI::ADE2 | Nakanishi et al., 2012 |
| pTSK548 | IAA-9Myc-KANMX cassette from | Morawska and Ulrich, 2013 |
| pTSK633 | pTEF-OsTIR1::URA3 | Kaur et al., 2018 |
| SGB14001 | pRS403-GFP-LacI::HIS3 modified at one of the *XhoI* sites flanking LacI gene, by Klenow treatment | This study |
| SGB14003 | pRS-SCM3-LacI::HIS3, PCR amplified SCM3 gene (with PA61-62 primers) inserted into vector backbone of SGB14001 in exchange of GFP gene constructing SCM3-LacI construct | This study |
| pSLB57 | pRS-Ask1-LacI::HIS3 | Lacefield et al., 2009 |

**Supplementary file 1C: List of primers used in ChIP-qPCR:**

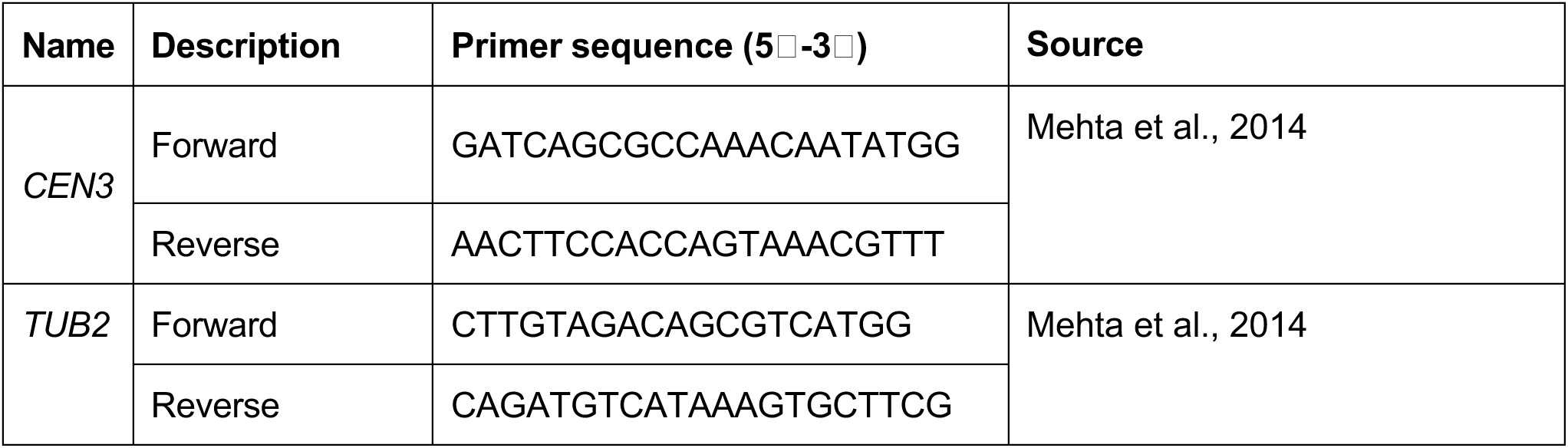

